# Molecular signature of pruriceptive MrgprA3^+^ neurons

**DOI:** 10.1101/727396

**Authors:** Yanyan Xing, Junyu Chen, Henry Hilley, Haley Steele, Jingjing Yang, Liang Han

**Affiliations:** School of Biological Sciences, Georgia Institute of Technology, Atlanta, GA 30332; Department of Human Genetics, Emory University School of Medicine, Atlanta, GA 30322

## Abstract

Itch, initiated by the activation of sensory neurons, is frequently associated with dermatological or systemic diseases and significantly affects patient quality of life. MrgprA3^+^ sensory neurons have been identified as one of the major itch-sensing neuronal populations. Mounting evidence has demonstrated that peripheral pathological conditions induce physiological regulations of sensory neurons, which is critical for the maintenance of chronic itch sensation. However, the underlying molecular mechanisms are not clear. Here we performed RNA sequencing of genetically labeled MrgprA3^+^ neurons under both naïve and allergic contact dermatitis condition. Our results revealed the unique molecular signature of itch-sensing neurons and the distinct transcriptional profile changes that result in response to dermatitis. We found enrichment of nine Mrgpr family members and two histamine receptors in MrgprA3^+^ neurons, suggesting that MrgprA3^+^ neurons are the main, direct neuronal target for histamine and Mrgprs agonists. In addition, Ptpn6 and Pcdh12 were identified as novel and highly selective markers of MrgprA3^+^ neurons. We also discovered that MrgprA3^+^ neurons respond to skin dermatitis in a way that is unique from other sensory neurons by regulating a combination of transcriptional factors, ion channels, and key molecules involved in synaptic transmission. These results significantly increase our knowledge of itch transmission and uncover potentially novel targets for combating itch.

## INTRODUCTION

Chronic itch is a devastating symptom frequently associated with dermatological diseases such as atopic dermatitis, psoriasis, and lichen planus (Yosipovitch and Bernhard, 2013). The significant discomfort associated with chronic itch not only leads to scratching, which worsens skin inflammation and causes secondary infection, but also influences the psychosocial status of the patients, triggering sleep disruption and further reducing quality of life (Leader et al., 2015).

Itch sensation is initiated by the activation of sensory neurons whose cell bodies reside in the dorsal root ganglia (DRG) or trigeminal ganglia (TG). Skin innervating sensory neurons project their peripheral and central axons to the skin and spinal cord respectively, providing a bridge for information flow from the peripheral skin to the central nervous system. Cutaneous stimuli or insults in the inflamed skin, such as pruritogens, are transduced into neuronal activities in the skin nerves to activate itch neural circuits and convey the signals to the brain (Ikoma et al., 2006).

Sensory neurons are highly diverse and have been classified into distinct groups based on their electrophysiological properties, expression of molecular markers and functional implications (Basbaum et al., 2009; Dong and Dong, 2018; Han and Dong, 2014; Zimmerman et al., 2014). Recent single-cell sequencing-based technologies revolutionized basic biomedical research and established a powerful and unbiased strategy for classification of neuronal types (Chiu et al., 2014; Hu et al., 2016; Kupari et al., 2019; Li et al., 2016; Usoskin et al., 2015). Three pruriceptive sensory neuron subtypes have been identified by both functional studies and single-cell RNAseq analysis and named NP1-3, each of which expresses distinct itch receptor combinations (Usoskin et al., 2015). NP1 neurons are classified by the expression of MrgprD (Liu et al., 2012), whereas NP2 neurons are classified by the expression of MrgprA3 and MrgprC11 (Han et al., 2013; Liu et al., 2009). Lpar3 and Lpar5 (Lysophosphatidic acid receptor 3 and 5) are highly expressed in NP1 population as well (Liu et al., 2012; Oude Elferink et al., 2015). NP3 neurons express the most itch receptors, including two co-receptors for IL-31 (Il31ra and osmr), Cysltr2 (cysteinyl leukotriene receptor 2), Htr1f (5-Hydroxytryptamine Receptor 1F) and S1pr1 (sphingosine-1-phosphate receptor 1) (Huang et al., 2018b; Mishra and Hoon, 2013; Solinski et al., 2019; Storan et al., 2015; Usoskin et al., 2015).

Peripheral pathological conditions can profoundly change the physiological properties of the innervating sensory neurons and modulate the communication of sensory neurons with second-order spinal neurons by utilizing both transcriptional and translational modifications (Basbaum et al., 2009; Piomelli and Sasso, 2014; Waxman and Zamponi, 2014). Indeed, ample evidence has demonstrated that nerve injury and inflammation can induce dysregulation of ion channel expression, which results in enhanced neuronal excitability that underlies chronic neuropathic and inflammatory pain (Waxman and Zamponi, 2014). This notion is further supported by the morphological and physiological changes of itch-sensing neurons under chronic itch conditions (Akiyama et al., 2010; Qu et al., 2014; Valtcheva et al., 2015; Zhu et al., 2017). For example, our recent studies demonstrated that NP2 MrgprA3^+^ neurons exhibit elevated itch receptor expression and hyperinnervation in the skin in a mouse dry skin model (Zhu et al., 2017) and enhanced excitability and spontaneous activities in a contact dermatitis model (Qu et al., 2014). Although the molecular mechanisms underlying those changes are not clear, RNA sequencing technology provides a reliable tool to reveal transcriptional level changes in the cells. Gene expression profiling of sensory neurons under a variety of pathological pain conditions have been investigated at either whole neuronal population or single-cell level (Aczel et al., 2018; Chung et al., 2016; Hu et al., 2016; Kathe and Moon, 2018; Starobova et al., 2019). However, analysis of itch-sensing neurons under chronic itch conditions have never been reported. Here we performed transcription profiling of MrgprA3^+^ neurons and investigated the change of its transcriptome following allergic contact dermatitis. The results will provide mechanistic insight into the molecular basis of chronic itch associated with atopic contact dermatitis and help to identify potential therapeutic targets for treatment.

## RESULTS

### MrgprA3^+^ neurons are a distinct subpopulation of DRG sensory neurons

In our previous study we generated a *MrgprA3*^*GFP-Cre*^ line, which allows genetic labeling of MrgprA3^+^ neurons (Han et al., 2013). We treated the lower back skin of *MrgprA3*^*GFP-Cre*^ mice with an allergen, squaric acid dibutylester (SADBE), to induce allergic contact dermatitis (ACD). We selected the lower back skin to minimize possible scratching-induced skin injury since mice are not able to scratch their lower back skin effectively. The treatment induced parakeratosis, dense inflammatory cell infiltration in the skin (Figure 1A), and spontaneous scratching behavior (Figure 1B), mimicking the symptoms of human patients with ACD. Thoracic and lumbar DRG (T11-L6) innervating the treated skin area were isolated after treatment and enzymatically dissociated. MrgprA3^+^ neurons and MrgprA3^−^ neurons were FACS purified based on GFP fluorescence, after DAPI staining to exclude dead cells, and the total RNA was extracted for RNA-seq analysis. cDNA libraries of MrgprA3^+^ and MrgprA3^−^ neurons were generated, sequenced, and analyzed for both the control and ACD treatment groups. Principal Components Analysis (PCA) showed clear segregation of MrgprA3^+^ and MrgprA3^−^neurons, both before and after the ACD treatment (Figure 1C), demonstrating that MrgprA3^+^ neurons are a distinct subpopulation of DRG sensory neurons.

**Figure 1.**
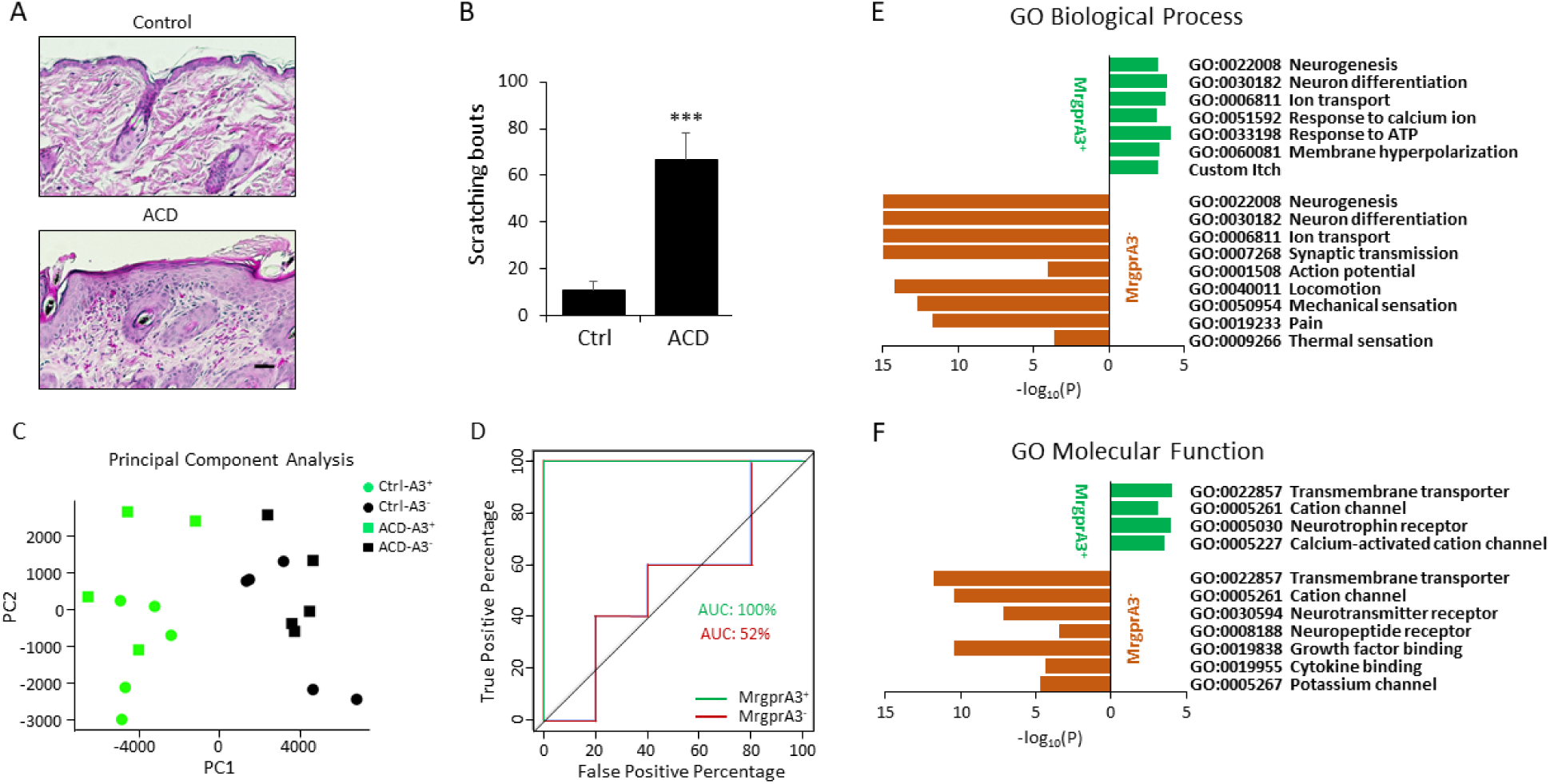
MrgprA3^+^ neurons are distinct itch-sensing neurons. (A) Hematoxylin and eosin (H&E) staining of the skin sections collected from *MrgprA3*^*GFP-Cre*^ mice after SADBE or vehicle control treatments. SADBE induced contact dermatitis in the skin evidenced by the parakeratosis and inflammatory cells infiltration in the skin. (B) *MrgprA3*^*GFP-Cre*^ mice developed robust scratching behavior after the allergic contact dermatitis treatment (C) Principal component analysis shows distinct transcriptome segregation of MrgprA3^+^ neurons and MrgprA3^−^ neurons. (D) Control and ACD conditions were classified with SVM-based classification model. The area under the ROC curve (AUC) is clearly higher for the MrgprA3^+^ neuron samples compare to MrgprA3^−^ neurons. (E-F) Gene ontology (GO) Biological Process (E) and Molecular Function (F) categories analysis of enriched genes in MrgprA3^+^ and MrgprA3^−^ neurons (see also Supplementary Table 2). All -log_10_(P) >15 was defined as 15 to fit the graph. \*\*\**P* < 0.005. Two-tailed unpaired Student’s t-test. Error bars represent s.e.m. Scale bar = 100 μm.

We further quantitatively evaluated how transcriptome profiles are different between control and ACD treatment groups within the MrgprA3^+^ and MrgprA3^−^ neuron samples. Particularly, we used the top 11 principal components of the transcriptome profiles (explaining 99% transcriptome variation) as features to fit a classification model to classify control and ACD treatments group (Cortes and Vapnik, 1995), respectively for MrgprA3^+^ and MrgprA3^−^ samples. As shown in the plot of the receiver operating characteristic (ROC) curve of the classification results (Fawcett, 2006) (Figure 1D), the area under the ROC curve (AUC) is clearly higher for the MrgprA3^+^ neuron samples, demonstrating better classification results. This data suggest that the MrgprA3^+^ neurons exhibited larger transcriptome profile change compare to the MrgprA3^−^ neurons after ACD treatment, consistent with the key role MrgprA3^+^ neurons play in mediating ACD itch.

### Transcriptional profile of MrgprA3^+^ neurons

We first compared the data from MrgprA3^+^ and MrgprA3^−^ populations in the control group to reveal the distinct transcriptional profile of MrgprA3^+^ neurons. In total 1,845 genes, including 584 upregulated genes and 1,261 downregulated genes, were found to be differentially expressed in MrgprA3^+^ neurons (Supplementary Table 1). We analyzed the specific Gene Ontology (GO) categories of the enriched genes in both neuronal populations (Figure 1E and 1F, Supplementary Table 2). Significant GO Biological Process terms of the genes enriched in MrgprA3^+^ neurons include itch (Custom GO Term), membrane hyperpolarization (GO:0060081), and response to ATP (GO:0033198), consistent with MrgprA3^+^ neurons function as itch-sensing neurons. Conversely, GO terms associated with genes enriched in MrgprA3^−^ neurons include pain (GO:0019233), thermal sensation (GO:0009266), mechanical sensation (GO:0050954), and locomotion (GO:0040011), which is unsurprising because MrgprA3^−^ neurons include pain sensors, touch sensors, and proprioceptors. GO terms such as neurogenesis (GO:0022008), neuron differentiation (GO:0030182), and ion transport (GO:0006811) are associated with genes enriched in both groups, suggesting that the two neuronal populations engage different gene sets to perform the same biological process. Similar results were also demonstrated by the GO Molecular Function analysis (GO:0022857 transmembrane transporter activity and GO:0005261 cation channel activity). Many GO categories associated with MrgprA3^+^ neurons are related to ion channels such as ion transport (GO:0006811), membrane hyperpolarization (GO:0060081) in GO-BP, and calcium-activated cation channel (GO:0005227) in GO-MF, suggesting that MrgprA3^+^ neurons utilize distinct groups of ion channels to perform neuronal functions.

We next examined the gene expression pattern of known genes that are important for sensory neuron functions. The first group we analyzed were Mrgpr (mas-related G protein-coupled receptor) family members (Meixiong and Dong, 2017) (Figure 2A). Consistent with previous studies from our group and others, MrgprB4, MrgprC11, and MrgprA1 are enriched in MrgprA3^+^ neurons (Han et al., 2013; Li et al., 2016; Lou et al., 2015; Meixiong et al., 2019). Five other Mrgprs with unknown functions including MrgprA2b, MrgprA4, MrgprA7, MrgprA8, and MrgprB5 are also highly enriched in MrgprA3^+^ neurons. MrgprD is downregulated since MrgprD^+^ and MrgprA3^+^ neurons are two separate populations (Liu et al., 2012; Usoskin et al., 2015). It is interesting to see nine members of the same family enriched in MrgprA3^+^ neurons, which only constitute 8% of the DRG sensory neurons. Among them, three (A3, C11, and A1) have been previously identified as itch receptors (Liu et al., 2012; Liu et al., 2009). Whether other enriched members are involved in itch sensation will be an interesting topic for future investigation.

**Figure 2.**
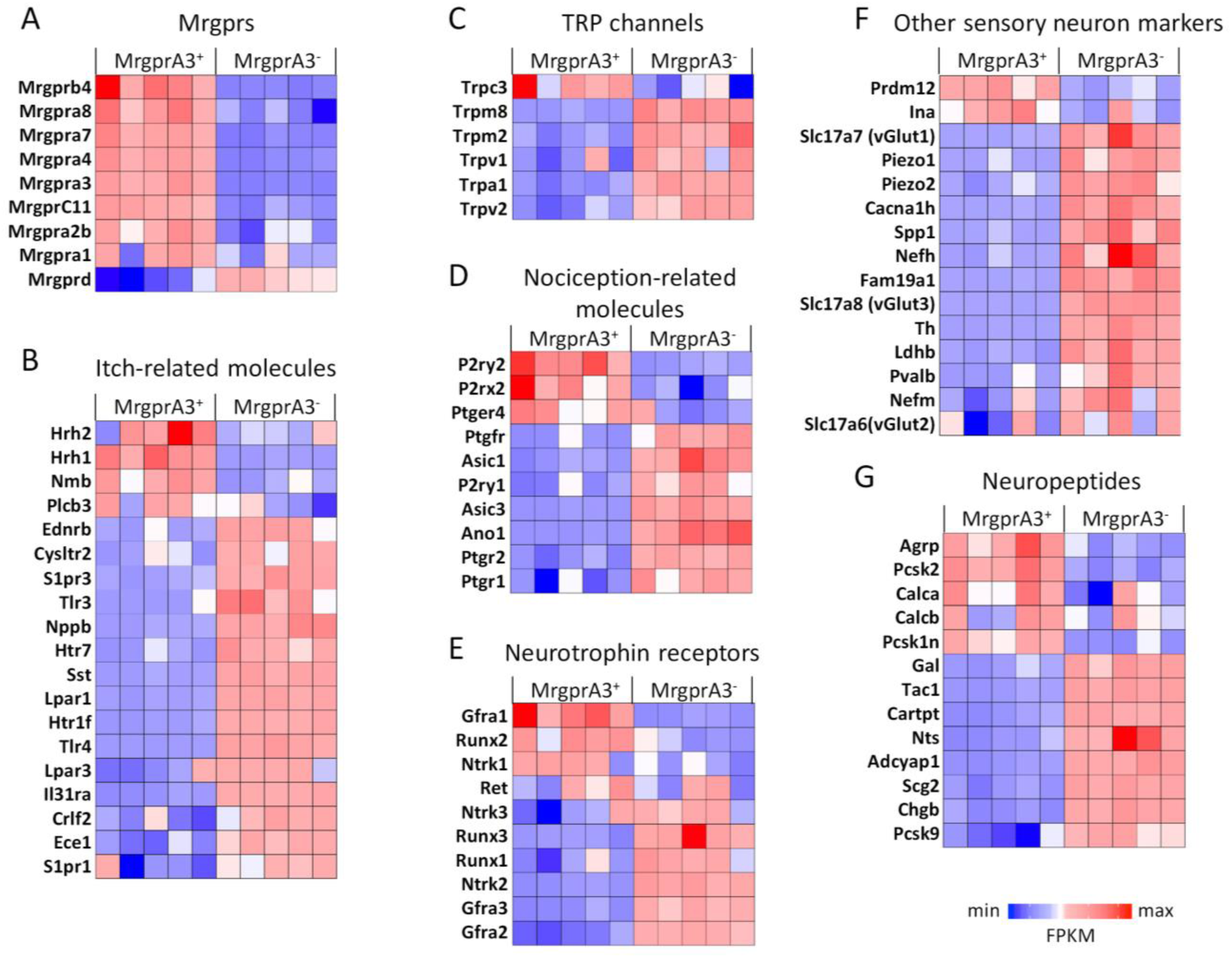
Heat-maps showing the expression pattern of sensory neuron markers in MrgprA3^+^ and MrgprA3^−^ neurons. Differentially expressed Mrgprs (A), itch-related molecules (B), TRP channels (C), nociception-related molecules (D), neurotrophin receptors (E), other sensory neuron markers (F), and neuropeptides (G) were analyzed and plotted in heat-maps. Columns are individual samples.

We then analyzed 19 known genes that play an important role in itch sensation (Figure 2B). Two histamine receptors (Hrh1 and Hrh2) and Plcb3, an isozyme mediating histamine-induced sensory neuron activation, are enriched in MrgprA3^+^ neurons (Han and Dong, 2014; Han et al., 2006). Neuromedian B (Nmb), a neuropeptide required for scratching behaviors induced by histamine, compound 48/80, and 5-HT (Wan et al., 2017), is also enriched in MrgprA3^+^ neurons. Nmb has been shown to be highly expressed in MrgprA3^+^ trigeminal sensory neurons innervating the conjunctiva and mediating ocular itch (Huang et al., 2018a). The enrichment of these genes and Mrgprs suggests that MrgprA3^+^ neurons are the primary, direct mediator of the Mrgprs agonists and histamine-induced itch.

As MrgprA3^−^ neurons include NP1 and NP3 populations (Usoskin et al., 2015), they are similarly enriched for many itch receptors and or itch-related molecules. The expression of Nppb (Natriuretic Peptide B) and Sst (somatostatin) defines the pruriceptive NP3 population that does not express MrgprA3 (Solinski et al., 2019; Usoskin et al., 2015). Cysltr2 (cysteinyl leukotriene receptor 2), Htr1f (5-Hydroxytryptamine Receptor 1F), and S1pr1 (sphingosine-1-phosphate receptor 1) are all enriched in Nppb/Sst^+^ NP3 neurons and mediate leukotriene C4, serotonin, and sphingosine-1-phosphate-induced itch sensation, respectively (Solinski et al., 2019). Both co-receptors for IL-31 (Il31ra and osmr) are highly expressed in both MrgprA3^+^ and MrgprA3^−^ neurons, although MrgprA3^−^ neurons are more enriched for Il31ra, suggesting that MrgprA3^+^ neurons are part of the neuronal population mediating Il-31-induced itch. Lysophosphatidic acid is a pruritogen in the sera of cholestatic patients with pruritus (Kremer et al., 2010). Lpar3 (Lysophosphatidic acid receptor 3), one of the cognate receptors for Lysophosphatidic acid, is exclusively expressed in pruriceptive NP1 population defined by the expression of MrgprD^+^ neurons and is downregulated in MrgprA3^+^ neurons. The other receptor examined for Lysophosphatidic acid (Lpar1) is also downregulated in MrgprA3^+^ neurons. The expression of both endothelin receptors and endothelin converting enzyme 1 (Ece1), which promotes endothelin receptor recycling and serves as a negative regulator of endothelin-induced itch (Kido-Nakahara et al., 2014), is relatively higher in MrgprA3^−^ population. Htr7 (5-Hydroxytryptamine Receptor 7), which couples to TRPA1 to mediate serotonergic itch (Morita et al., 2015), is preferentially expressed in MrgprA3^−^ neurons. The expression of some receptors such as Protease-activated receptor 2 (Par2) is not clear in our RNAseq analysis probably due to unreliable detection.

Transient receptor potential (TRP) ion channels are key sensors for heat, cold, and reactive chemicals in sensory neurons (Basbaum et al., 2009). They are often recruited by G protein-coupled receptors to induce calcium influx and action potential generation to respond to noxious or pruriceptive stimuli. For example, histamine receptor 1 (Hrh1) activates TRPV1 to induce neuronal activation (Shim et al., 2007), whereas MrgprA3 and MrgprC11 recruit TRPA1 (Wilson et al., 2011). Although both TRPA1 and TRPV1 are highly expressed in MrgprA3^+^ neurons, they both are expressed in a very large population of small-diameter neurons. Thus, the present analysis demonstrates the enrichment of TRPA1 and TRPV1 in MrgprA3^−^ neurons. Cold sensory TRPM8 is significantly enriched in MrgprA3^−^ neurons. The only TRP channel enriched in MrgprA3^+^ neurons is TRPC3, whose function in itch sensation is not yet clear (Figure 2C).

We next examined the expression patterns of several families that are key players in nociception, which include purinergic receptors, prostaglandin receptors, and acid-sensing ion channels (Basbaum et al., 2009) (Figure 2D). Three genes are observed to be specifically enriched in MrgprA3^+^ neurons including P2rx2 (ionotropic purinergic receptor 2), P2ry2 (metabotropic purinergic receptor 2), and Ptger4 (prostaglandin E receptor 4, or EP4). The enrichments of P2rx2 and P2ry2 raise the question if ATP plays a role in itch induction, which has not been intensively investigated yet. Previous studies have shown an increased level of prostaglandin E2, an agonist for EP4, in the skin of patients with atopic dermatitis (Fogh et al., 1989). Consistently, blockade of PGE2 signaling reduced spontaneous scratching behavior in a mouse model of dermatitis (Emrick et al., 2018). Our analysis suggests that PGE2 might activate MrgprA3^+^ neurons via the activation of EP4.

In the ‘Other sensory neuron markers’ category (Figure 2F), as expected, many markers for non-pruriceptive neurons are downregulated in MrgprA3^+^ neurons. Markers for large-diameter neurons (vGlut1, Ldhb, Cavna1h, Spp1, and Fam19a1), markers for proprioceptors (Pvalb), and marker for C-low-threshold mechanoreceptors (TH and vGlut3) (Usoskin et al., 2015) are mainly enriched in the MrgprA3^−^ population. vGlut2 is highly expressed in both populations. Both Piezo1 and Piezo2 are enriched in MrgprA3^−^ neurons. Prdm12 (PR Domain Zinc Finger Protein 12), a transcriptional regulator controlling the generation of nociceptive lineage during development, is enriched in MrgprA3^+^ neurons (Bartesaghi et al., 2019; Desiderio et al., 2019). Different types of neurofilaments are expressed by a distinct subset of sensory neurons as their major cytoskeleton components. Nefm (neurofilament medium chain) and Nefh (neurofilament heavy chain) are enriched in MrgprA3^−^ neurons. By contrast, Ina, an internexin neuronal intermediate filament, is enriched in MrgprA3^+^ neurons. Expression of neurotrophin receptors has long been one of the criteria to classify somatosensory neurons (Lallemend and Ernfors, 2012). MrgprA3^+^ neurons are enriched for Gfra1 (GDNF receptor alpha 1) and Ntrk1 (TrkA, neurotrophic receptor tyrosine kinase 1). Interestingly, we also observed the enrichment of Runx2 (runt related transcription factor 2), the only Runx family member that has not been linked to somatosensory neurons differentiation and function, in MrgprA3^+^ neurons (Figure 2E).

Neuropeptides are one of the major types of molecules mediating neuronal transduction. In the somatosensory neural pathway, several neuropeptides have been demonstrated to be neurotransmitters or neuromodulators, such as Nmb, GRP, Nppb, and Sst (Huang et al., 2018b; Mishra and Hoon, 2015; Solinski et al., 2019; Wan et al., 2017). Similar to the previous single-cell RNA-seq analysis, we found MrgprA3^+^ neurons are enriched for Nmb (neuromedin B), CGRP (Calca, and Calcb, calcitonin/calcitonin-related polypeptide, alpha and beta), and Agrp (Agouti-related peptide). In addition, we found that MrgprA3^+^ neurons are enriched for Pcsk2 (Proprotein Convertase Subtilisin/Kexin Type 2), and Pcsk1n (Proprotein Convertase Subtilisin/Kexin Type 1 Inhibitor), both of which are involved in the processing of neuropeptide and have never been linked to sensory neuron function (Figure 2G).

We next focused our analysis on the expression patterns of voltage-gated ion channels, which determine the generation of action potentials and neuronal firing patterns (Figure 3). We found that MrgprA3^+^ neurons employ unique combinations of ion channels to maintain the resting membrane potential and initiate the neuronal firing. For sodium channels (Figure 3A), Nav1.9 (Scn11a) is enriched in MrgprA3^+^ neurons, which is in line with its known role in itch sensation (Salvatierra et al., 2018). The other two enriched members in MrgprA3^+^ neurons are β subunits of voltage-gated sodium channels Scn2b and Scn3b, both of which have been linked based on genetic analysis to heart diseases such as Brugada Syndrome and atrial fibrillation (Dehghani-Samani et al., 2019), with unclear function in somatosensation. Scn2b has been shown to modulate tetrodotoxin-sensitive channel expression and activities, and has been linked to the development of mechanical allodynia (Lopez-Santiago et al., 2006). Besides, Cacng2 is the only enriched voltage-gated calcium channel in the MrgprA3^+^ neurons (Figure 3B). In spinal dorsal horn neurons, Cacng2 acts as an AMPA receptor regulatory protein and contributes to inflammatory pain (Sullivan et al., 2017; Tao et al., 2006). Its polymorphisms have been associated with chronic pain after mastectomy as well (Bortsov et al., 2019), although its role in itch transmission is not yet clear. Ttyh2 (Drosophila tweety homolog 2), a calcium-activated chloride channel (Suzuki, 2006), is highly enriched in MrgprA3^+^ neurons (Figure 3C). Potassium channels constitute the largest ion channel family and have been demonstrated to be differentially expressed across somatosensory subsets (Chiu et al., 2014; Usoskin et al., 2015). We observed seven potassium channels enriched in MrgprA3^+^ neurons with Kcnk18 as the only enriched leaky potassium channel (TRESK, potassium channel, subfamily K, member 18) (Figure 3D) (Sano et al., 2003).

**Figure 3.**
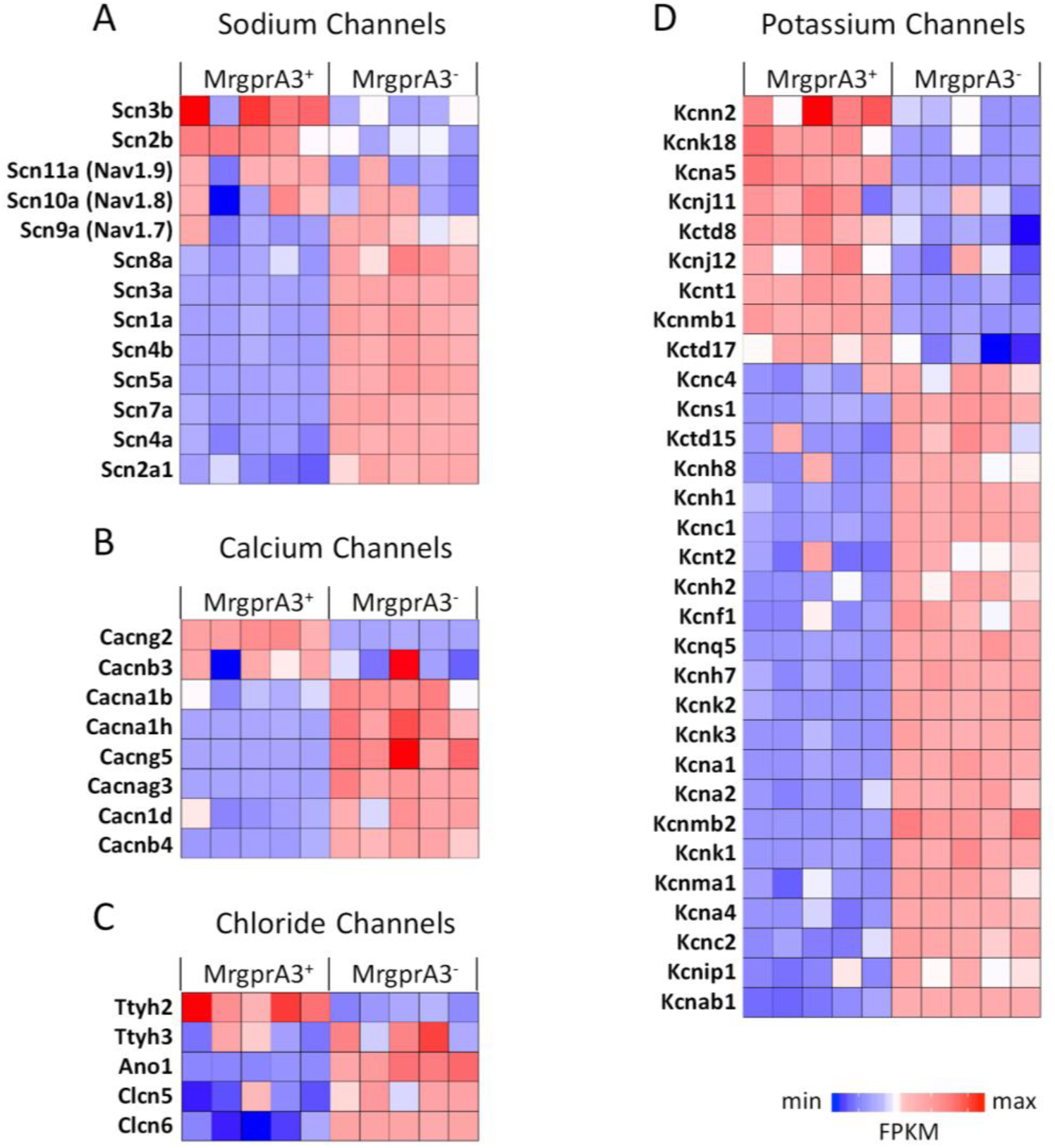
Heat-maps showing the expression pattern of ion channels. Differentially expressed sodium channels (A), calcium channels (B), chloride channels (C), and potassium channels (D) were analyzed and plotted in heat-maps. Columns are individual samples.

To validate the enriched transcripts *in vivo*, we chose two novel genes identified to be highly enriched in MrgprA3^+^ neurons for RNAscope *in situ* hybridization analysis (Figure 4). Ptpn6 (Protein Tyrosine Phosphatase Non-Receptor Type 6) is expressed in 3.4% (96/2830) of DRG sensory neurons (Figure 4A). Double fluorescent *in situ* hybridization results confirm that 83.3% (80/96) of Ptpn6^+^ neurons are expressing MrgprA3. Pcdh12 (protocadherin 12) is expressed in 6.4% (133/2066) of DRG sensory neurons and the majority of Pcdh12^+^ neurons (69.9%, 93/133) coexpress MrgprA3. These results show that the enriched genes identified by our RNA-seq analysis also exist *in vivo*.

**Figure 4.**
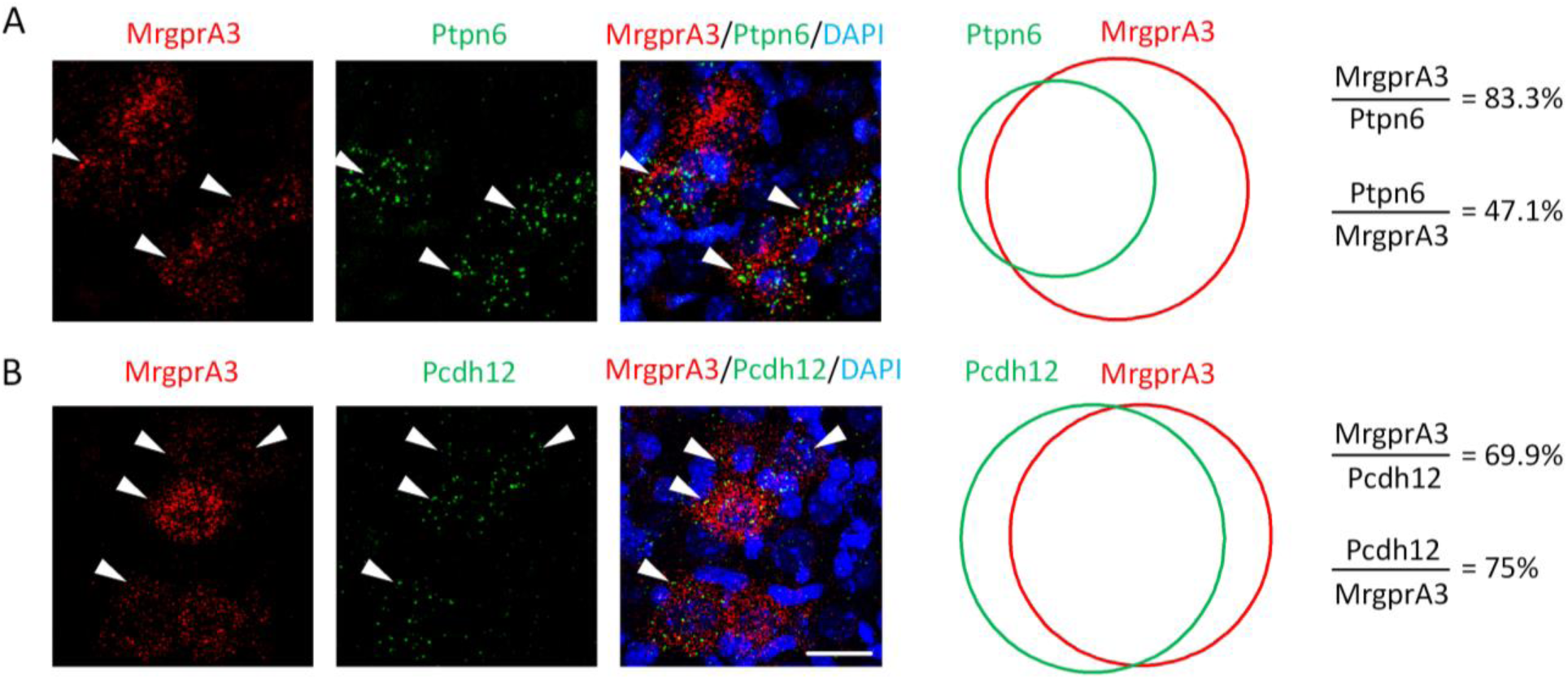
*In situ* hybridization validation of identified novel markers in MrgprA3^+^ neurons. (A-B) Fluorescent *in situ* hybridization and Venn diagrams showing the expression overlapping of Ptpn6 (A) and Pcdh12 (B) with MrgprA3. Scale bar = 20 μm.

### Transcriptional profile of MrgprA3^+^ neurons was changed by atopic contact dermatitis

It is well recognized that pathological skin conditions can profoundly change the innervating sensory neurons morphologically and functionally. To examine how dermatitis changes the expression profile of itch-sensing neurons, we compared the sequencing data from neurons isolated under control and ACD conditions. The results showed that dermatitis treatment significantly changed the expression profile of both MrgprA3^+^ neurons and MrgprA3^−^ neurons, demonstrating the common neuro-immuno crosstalk between inflamed skin and innervating sensory neurons. We have identified 408 upregulated genes and 425 downregulated genes in MrgprA3^+^ neurons. Similarly, 292 upregulated genes and 420 downregulated genes were identified in MrgprA3^−^ neurons (Figure 5A, Supplementary Table 3). Inflamed skin caused significant damage to all of the innervating sensory neurons, as demonstrated by the enrichment of GO terms such as programmed cell death (GO:0012501) and cellular response to stress (GO:0033554) in the differentially regulated genes in both types of neurons. Specific GO terms such as chromosome organization (GO:0051276) and histone modification (GO:0016570) are also associated with both groups, suggesting that skin inflammation affects the gene expression process in all innervating sensory neurons. However, the genes affected in the two neuronal populations are largely non-overlapping (Figure 5A), which indicates that MrgprA3^+^ neurons respond to the dermatitis condition in a very different way from other sensory neuron subtypes. Indeed, unique GO terms are identified to be associated with the two neuronal populations. GO terms such as regulation of catabolic process (GO:0009894), golgi vesicle transport (GO:0048193), and cellular response to DNA damage (GO:0006974) are unique to MrgprA3^+^ neurons, while chemical synaptic transmission (GO:0007268), cation transmembrane transport (GO:0098655), and MAPK (GO:0000165) are uniquely associated with MrgprA3^−^ neurons (Figure 5B).

**Figure 5.**
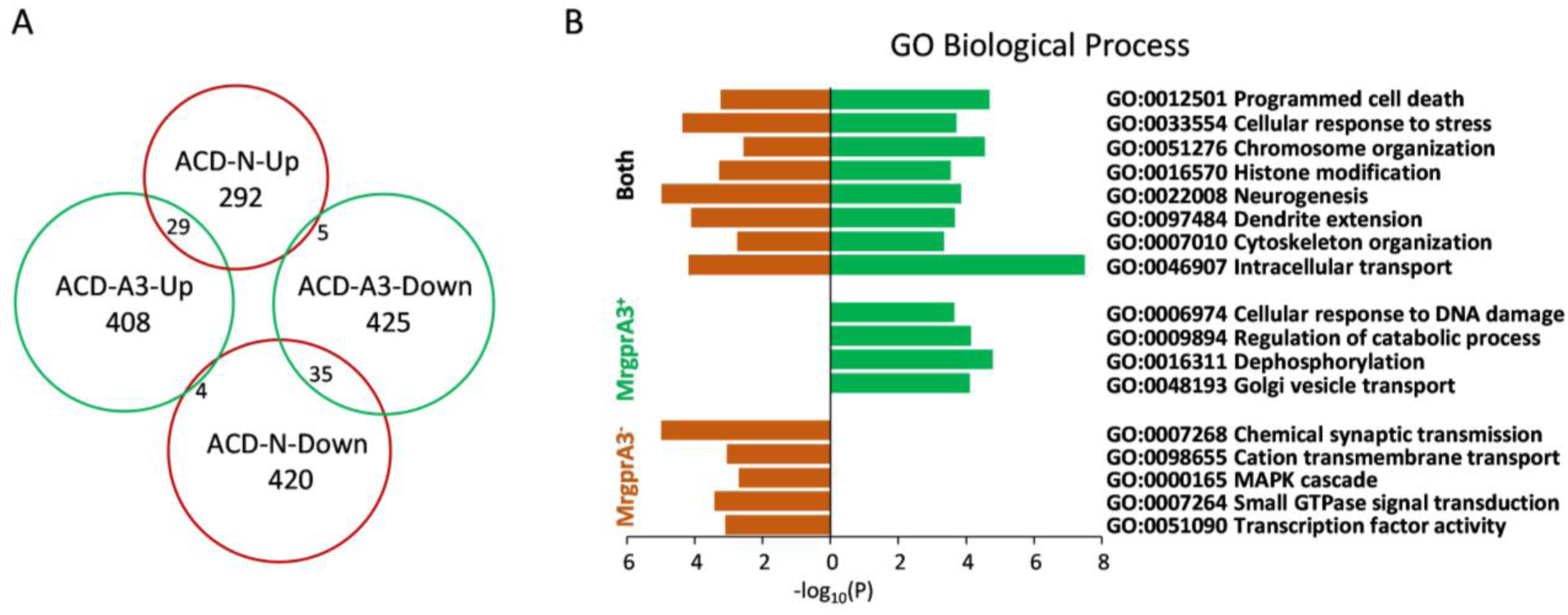
MrgprA3^+^ and MrgprA3^−^ neurons are differentially affected by the allergic contact dermatitis condition. (A) Venn diagram showing the minimal overlap of genes changed by ACD condition in MrgprA3^+^ and MrgprA3^−^ neurons. Numbers inside of the circle indicate the number of genes identified in the analysis. A3: MrgprA3^+^ neurons, N: MrgprA3^−^ neurons, ACD: allergic contact dermatitis. (B) Gene ontology analysis of genes changed by the ACD treatment (see also Supplementary Table 2).

A small percentage of affected genes (73 total in both Up and Down genes, Figure 5A, Supplementary Table 3) were changed in both neuronal populations, suggesting a common gene set might exist for sensory neurons to respond to the inflamed peripheral skin. Among them, 29 were upregulated in both neuronal populations, 35 were downregulated in both, and 9 showed different regulation in the two types of neurons. Although statistically significant GO Terms associated with those genes were not found, we identified several interesting genes including those related to inflammatory responses such as Il17ra (interleukin 17 receptor A), Tnfrsf1a (tumor necrosis factor receptor superfamily, member 1a), and Ptgfr (prostaglandin F receptor); those related to neuronal function such as Syt2 (synaptotagmin II) and Slc12a2 (Sodium/Potassium/Chloride Transporter, Solute Carrier Family 12, Member 2); and those related to intracellular signaling transduction in sensory neurons such as Pla2r1 (phospholipase A2 receptor 1) and Dgkz (diacylglycerol kinase zeta).

We next focused on gene transcripts that might contribute to the morphological or functional changes of sensory neurons. We first examined key transcription factors that are critical for changing the transcriptional profile. Distinct sets of transcription factors were identified within the two neuronal populations (Figure 6A), the majority of which lack known sensory neuron function. Etv5, an essential molecule regulating sensory neuron differentiation and axonal growth during development (Fontanet et al., 2013), is upregulated in MrgprA3^+^ neurons. Six4, a transcriptional factor required for the survival of sensory neurons, is downregulated in MrgprA3^+^ neurons (Konishi et al., 2006; Moody and LaMantia, 2015; Yajima et al., 2014). Runx1 and Prdm12, both of which are required for nociceptor specification, are downregulated in MrgprA3^−^ neurons.

**Figure 6.**
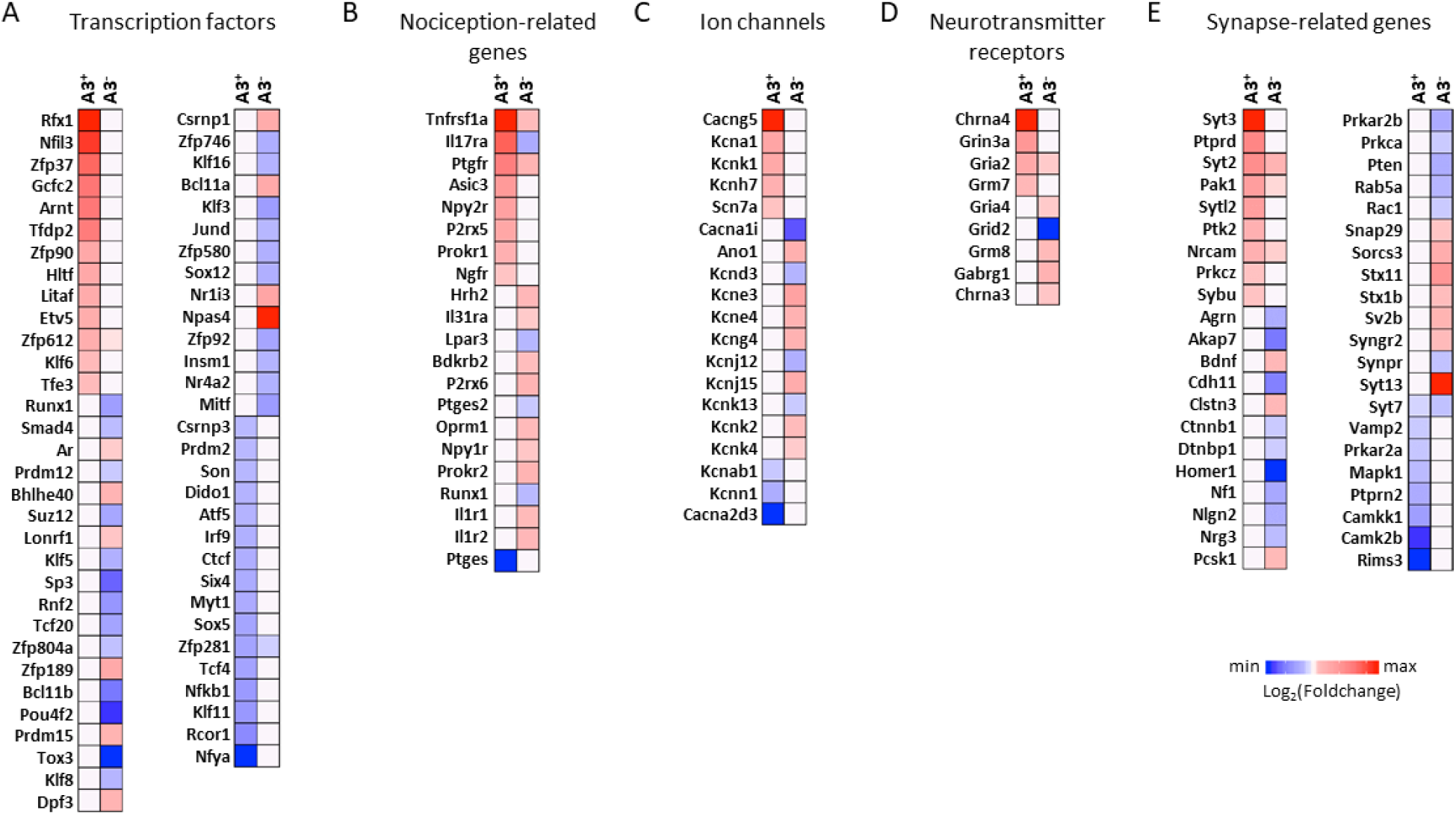
Heat-maps showing transcripts changed by ACD treatment in MrgprA3^+^ and MrgprA3^−^ neurons. Lists of transcription factors (A), nociception-related genes (B), ion channels (C), neurotransmitter receptors (D), synapse-related genes (E) whose expression were changed by the ACD treatment in MrgprA3^+^ and MrgprA3^−^ neurons. For genes that were not significantly changed by ACD (p>0.05), the value of log_2_(Foldchange) was defined as 0.

We then focused our analysis on the genes related to sensory neuron function (Figure 6B). Both MrgprA3^+^ and MrgprA3^−^ neurons showed distinct gene change pattern after ACD treatment. Three itch receptors (Hrh2, Il31ra, and Lpar3) were changed in MrgprA3^−^ neurons, while we did not observe any significant change of itch receptors in MrgprA3^+^ neurons. ASIC3 (acid-sensing ion channel 3), P2rx5 (ionotropic purinergic receptor 5), and Ptges (prostaglandin E synthase) were changed in MrgprA3^+^ neurons, while Bdkrb2 (Bradykinin receptor beta 2), P2rx6 (metabotropic purinergic receptor 6) and Ptges2 (prostaglandin E synthase 2) were changed in MrgprA3^−^ neurons. A TNF receptor (Tnfrsf1a) and several interleukin receptors (Il1r1, Il1ra, and Il17ra) were changed, indicating their involvement in sensory neurons in response to skin inflammation. Interestingly, we observed a significant increase of Ngfr (nerve growth factor receptor, or P75) only in MrgprA3^+^ neurons, consistent with the role of NGF in promoting chronic itch and sensory nerve sensitization (Ikoma et al., 2006; Mollanazar et al., 2016). Many ion channels have been identified as well, indicating the change of electrophysiological properties of neurons (Figure 6C). Among them, Cacng5 (calcium channel, voltage-dependent, gamma subunit 5) shows the greatest changes in MrgprA3^+^ neurons and Kcne3 (potassium voltage-gated channel, Isk-related subfamily, gene 3) shows the greatest change in MrgprA3^−^ neurons.

Studies have shown that peripheral insults can lead to changes in the sensory signal transmission to spinal cord neurons and presynaptic regulation by the local spinal circuits or descending fibers. Indeed, we found significant changes in many neurotransmitter receptors (Figure 6D). Among them, Gria2 (GluR2) was increased in both MrgprA3^+^ and MrgprA3^−^ neurons. Many genes related to synaptic transmission were also identified (Figure 6E-F) including Syt2 (synaptotagmin II), which was increased in both neuron types, and Syt7 (synaptotagmin VII), which was decreased in both neuron types.

## DISCUSSION

Molecular characterization of somatosensory neurons subtypes has been an effective approach to understanding sensory neuron function and circuitry. Although recent single-cell RNAseq analysis has revealed invaluable information on different types of sensory neurons (Chiu et al., 2014; Li et al., 2016; Usoskin et al., 2015), the detail molecular characteristics of itch-sensing neurons have not previously been extensively investigated. Here we present transcriptional profiling of pruriceptive MrgprA3^+^ neurons at the population level and reveal the unique molecular signature of those neurons. Moreover, we, for the first time, demonstrate distinct transcriptional profile changes in itch-sensing neurons that result from atopic contact dermatitis.

Mrgprs are a large G protein-coupled receptor family containing more than 20 coding members in mice, with many exclusively expressed in sensory neurons (Dong et al., 2001). Four Mrgprs including MrgprA1, MrgprA3, MrgprC11, and MrgprD have been identified as itch receptors and serve as molecular markers to classify pruriceptive neuronal populations (Liu et al., 2012; Liu et al., 2009; Meixiong et al., 2019). Interestingly, we found that MrgprA3^+^ neurons are enriched for eight other Mrgpr family members including two identified itch receptors MrgprA1 and MrgprC11, and five other members: MrgprB4, MrgprA2b, MrgprA4, MrgprA7, and MrgprA8. Two histamine receptors (Hrh1 and Hrh2) and Plcβ3 are also enriched in MrgprA3^+^ neurons. These results indicate that itch induced by histamine and Mrgprs agonists are primarily mediated by MrgprA3^+^ neurons.

Analogous to the T2Rs bitter receptor family in the gustation system and odorant receptor family in the olfaction system (Secundo et al., 2014; Yarmolinsky et al., 2009), Mrgpr family contains a large group of G protein-coupled receptors with different members detecting different chemical stimuli. MrgprA1 is activated by bilirubin and substance P and mediates cholestatic itch (Azimi et al., 2017; Meixiong et al., 2019). MrgprA3 responds to anti-malaria drug chloroquine (Liu et al., 2009). MrgprC11 mediate Bam8-22, SLIGRL, cathepsin S, and cysteine protease Der p1-induced itch (Liu et al., 2009; Liu et al., 2011; Reddy and Lerner, 2017; Reddy et al., 2015), while MrgprD is responsible for β-alanine-induced itch (Liu et al., 2012). In the olfactory system, each olfactory receptor neuron within the olfactory epithelium only expresses a single odorant receptor. The identity of odorants is encoded by a combinatorial receptor code with each odorant receptor responding to multiple odorants and each odorant activating multiple receptors (Secundo et al., 2014). This strategy allows the olfactory system to distinguish a massive number of odorant molecules, which is critical for recognizing food, predators, and mates. In contrast, in gustation system, it is more important for bitter taste to reveal the safety of potential food than to identify different bitter compounds. Therefore, multiple bitter taste receptors are co-expressed in the same taste receptor cells and stimulate the same neural circuits upon activation (Yarmolinsky et al., 2009). Our results demonstrate that the somatosensory system employs the same strategy as gustation system for detecting itch. Multiple Mrgpr itch receptors are highly enriched in MrgprA3^+^ neurons to ensure the detection of the somatosensory modality of the stimuli (itchy), while the identity of each pruritogen, which is not critical for the safety and survival of the animals, can be ignored. Similarly, the pruriceptive NP3 neurons also express multiple itch receptors including Il31ra, osmr, Cysltr2, Htr1f, and S1pr1. This expression pattern of itch receptors supports the hypothesis that itch sensing is mediated by a labeled line in the peripheral sensory neurons (Han and Dong, 2014).

Our sequencing analysis reveals the expression of unique combinations of GPCRs, neuropeptides, neurotrophic factors, and ion channels in MrgprA3^+^ neurons. The analysis of many genes is consistent with previous single-cell RNA-seq analysis such as the enrichment of MrgprB4, Rspo1, and Gfra1 in MrgprA3^+^ neurons and the enrichment of Il31ra, Nppb, Sst, and Lpar3 in MrgprA3^−^ neurons (Li et al., 2016; Usoskin et al., 2015). Our analysis also discovered several genes that have not previously been identified in MrgprA3^+^ neurons such as MrgprA4, MrgprA7, Ptpn6, and Pcdh12. Understanding the function of those genes in itch transmission will possibly lead to a better understanding of the basic mechanisms of itch transmission.

In skin, there are numerous cellular contacts between all innervating sensory nerves with cutaneous cells and immune cells. Therefore, it is not a surprise to observe significant changes in the expression profile of both MrgprA3^+^ neurons and MrgprA3^−^ neurons within atopic contact dermatitis. However, PCA-AUC analysis demonstrated that the ACD condition generated a much larger effect on MrgprA3^+^ neurons than it did on MrgprA3^−^ neurons, emphasizing the critical role of MrgprA3^+^ neurons in mediating ACD itch. Moreover, the genes affected in the two neuronal populations are largely non-overlapping, demonstrating the distinct gene regulation pattern of MrgprA3^+^ neurons by the ACD condition.

Chronic itch is induced by the generation of a high level of endogenous agonists for itch-sensing neurons in pathological conditions. It can be enhanced and maintained by the abnormal function of sensory neurons, as well as changes in the synaptic transmission between sensory neurons and spinal neurons. We did not observe any itch receptor change in MrgprA3^+^ neurons, suggesting that skin inflammation does not regulate receptor expression to changes neuronal sensitivity. However, we observed a change in the expression of genes in the following categories: transcriptional factors (such as Etv5 and Six4), purinergic receptor (such as P2rx5), ion channels (such as ASIC3 and Cacng5), neurotransmitter receptors (Gria2), and key molecules in synaptic transmission (such as Syt2 and Syt7), all of which may contribute to the generation of chronic itch in dermatitis skin. The change in the TNF receptor (Tnfrsf1a) and interleukin receptors (Il1r1, Il1ra, and Il17ra) also suggest one possible way that the sensory nerves respond to the inflammatory condition in the skin. In MrgprA3^−^ neurons, we observed the upregulation of three itch receptors Hrh2, Il31ra, and Lpar3 after ACD treatment. Lpar3 and Il31ra are highly expressed in pruriceptive NP1 and NP3 population respectively. These results suggest the involvement of all three types of pruriceptive sensory neurons in mediating ACD itch.

In summary, our study reveals the distinct transcriptional profile of pruriceptive MrgprA3^+^ neurons and unique gene regulation in response to peripheral allergic inflammation. Further functional studies investigating the role of newly identified genes in chronic itch will provide novel insights into the basic molecular and cellular mechanisms of itch and help to discover novel therapeutic targets for relieving itch.

## MATERIALS AND METHODS

### Mice

All experiments were performed with approval from the Georgia Institute of Technology Animal Use and Care Committee. The MrgprA3-GFP-Cre line was generated previously and bred in the animal facility at Georgia Institute of Technology. All the mice used had been backcrossed to C57BL/6 mice for at least ten generations. 2-to 3-month old males (20 - 30g) were used for contact dermatitis treatment. Mice were housed in the vivarium with 12-hr-light/dark cycle. The housing group was 5 at maximum with food and water ad libitum.

### Contact dermatitis treatment

A mouse model of allergic contact dermatitis is produced by treating the skin with an allergen squaric acid dibutylester (SADBE, 25ul, 0.5% in acetone, Sigma-Aldrich, St. Louis, MO). On day 1-3, SADBE was applied to the shaved abdomen once a day to initiate T lymphocyte sensitization. On day 8-12, SADBE was applied to the shaved lower back skin to induce dermal inflammation. Since mice are not able to scratch their lower back efficiently, this skin area was selected for treatment to minimize scratching-induced skin injury. On day 13, 8 pairs of DRG in the lower thoracic region and lumbar region (T11-L6) that innervate the treated skin area were dissected for FACS sorting and RNA isolation. Animals were recorded for 60 min to examine spontaneous scratching behaviors before DRG dissection.

### Flow cytometry and RNA isolation

DRG were collected in cold DH10 medium and digested in Collagenase/Dispase solution (Roche, Indianapolis, IN) at 37 °C for 30 min. After trituration, cells were filtered through a 100 μm strainer, centrifuged, re-suspended in HBSS and subjected to flow cytometry. GFP^+^ cells and GFP^−^ cells were separated into different Eppendorf tubes containing RNAlater to preserve RNA quality at the time of isolation. Total RNA was extracted using the RNeasy Micro Kits with on column genomic DNA digestion (Qiagen, Germantown, MD) according to the manufacturer’s protocol.

### RNA-seq and bioinformatics analysis

Library construction and RNA-seq were performed by BGI Genomics. Briefly, RNA quality was determined by Agilent 2100 Bioanalyzer (Agilent Technologies, Santa Clara, CA). Samples with RIN >7 were used for analysis. Polyadenylated mRNA libraries were generated using the oligo(dT) magnetic beads and were used for cDNA library construction, amplification and sequencing. RNA-seq was performed with the illumina HiSeq™ 4000 at a depth of 50 million single-end reads of 50 bp. After QC, the raw reads were filtered by removing adaptor sequences, sequences with more than 10% of unknown bases, and low-quality bases. After filtering, clean reads are mapped to reference genome by Bowtie2 (Langmead and Salzberg, 2012), and then mapped read counts were normalized to the number of Fragments Per Kilobase of transcript per Million mapped reads (FPKM). Principal component analysis (PCA) was conducted using the FPKM data. To classify ACD and Ctrl conditions, we used support vector machine (svm) method (Cortes and Vapnik, 1995; Fawcett, 2006) to analyze the MrgprA3^+^ and MrgprA3^−^ samples, taking the top 11 PCs that explain 99% of the transcriptome profile variance. The results suggest the transcription profiles can distinguish ACD from Ctrl conditions better within MrgprA3^+^ samples than MrgprA3^−^ samples. We used the DESeq2 tool (Love et al., 2014)to identify differentially expressed genes (DEGs). Criteria to identify DEGs in Ctrl-MrgprA3^+^ vs Ctrl-MrgprA3^−^ neurons comparison is FDR < 0.05 and fold change > 1.85. Criteria to identify DEGs in Ctrl-MrgprA3^+^ vs ACD-MrgprA3^+^ comparison is p < 0.05 and fold change > 1.3. All DEGs (totally 584) enriched in MrgprA3^+^ neurons and top 700 DEGs enriched in MrgprA3^−^ neurons were identified and subjected for gene ontology (GO) analysis using WEB-based GEne SeT AnaLysis Toolkit (WebGestalt). All DEGs (833 genes in MrgprA3^+^ neurons and 712 genes in MrgprA3^−^ neurons) in Ctrl-MrgprA3^+^ vs AEW-MrgprA3^−^ neurons comparison were used for ontology analysis. Heat-maps were generated using the normalized read counts per gene. To enhance the visibility of the heat-maps, the read counts were normalized by subtracting the median count per gene and divided by the median absolute deviation per gene.

### *In situ* hybridization

Lumbar DRGs (L3-L6) were dissected from WT C57BL/6J mice. In situ hybridization was performed using the RNAscope fluorescent multiplex kit (ACD Cat#320850) according to the manufacturer’s instructions. Probes used: MrgprA3-C2 (ACD 548161-C2), Ptpn6 (ACD 450081), Pcdh12 (ACD 489891), Rspo1 (ACD 401991), and Mustn1 (CAD 568751). Images were collected on a Zeiss LSM700 confocal microscope system.

## DATA AVAILABILITY

The gene data sets will be deposited to NCBI Gene Expression Omnibus after paper acceptation.

## ACKNOWLEDGMENTS

We thank the Physiological Research laboratory (PRL) at Georgia Institute of Technology for the animal care and services. The work was supported by grants from the US National Institutes of Health to L.H. (NS087088 and HL141269), Pfizer Aspire Dermatology Award to L.H. and start-up funding from the Department of Human Genetics at Emory University to J.Y.

## References

Aczel, T., Kun, J., Szoke, E., Rauch, T., Junttila, S., Gyenesei, A., Bolcskei, K., and Helyes, Z. (2018). Transcriptional Alterations in the Trigeminal Ganglia, Nucleus and Peripheral Blood Mononuclear Cells in a Rat Orofacial Pain Model. Frontiers in molecular neuroscience 11, 219.

Akiyama, T., Carstens, M.I., and Carstens, E. (2010). Enhanced scratching evoked by PAR-2 agonist and 5-HT but not histamine in a mouse model of chronic dry skin itch. Pain 151, 378–383.

Azimi, E., Reddy, V.B., Pereira, P.J.S., Talbot, S., Woolf, C.J., and Lerner, E.A. (2017). Substance P activates Mas-related G protein-coupled receptors to induce itch. J Allergy Clin Immunol 140, 447–453 e443.

Bartesaghi, L., Wang, Y., Fontanet, P., Wanderoy, S., Berger, F., Wu, H., Akkuratova, N., Boucanova, F., Medard, J.J., Petitpre, C., et al. (2019). PRDM12 Is Required for Initiation of the Nociceptive Neuron Lineage during Neurogenesis. Cell Rep 26, 3484–3492 e3484.

Basbaum, A.I., Bautista, D.M., Scherrer, G., and Julius, D. (2009). Cellular and molecular mechanisms of pain. Cell 139, 267–284.

Bortsov, A.V., Devor, M., Kaunisto, M.A., Kalso, E., Brufsky, A., Kehlet, H., Aasvang, E., Bittner, R., Diatchenko, L., and Belfer, I. (2019). CACNG2 polymorphisms associate with chronic pain after mastectomy. Pain 160, 561–568.

Chiu, I.M., Barrett, L.B., Williams, E.K., Strochlic, D.E., Lee, S., Weyer, A.D., Lou, S., Bryman, G.S., Roberson, D.P., Ghasemlou, N., et al. (2014). Transcriptional profiling at whole population and single cell levels reveals somatosensory neuron molecular diversity. Elife 3.

Chung, M.K., Park, J., Asgar, J., and Ro, J.Y. (2016). Transcriptome analysis of trigeminal ganglia following masseter muscle inflammation in rats. Molecular pain 12.

Cortes, C., and Vapnik, V. (1995). Support-Vector Networks. Mach Learn 20, 273–297.

Dehghani-Samani, A., Madreseh-Ghahfarokhi, S., and Dehghani-Samani, A. (2019). Mutations of Voltage-Gated Ionic Channels and Risk of Severe Cardiac Arrhythmias. Acta Cardiol Sin 35, 99–110.

Desiderio, S., Vermeiren, S., Van Campenhout, C., Kricha, S., Malki, E., Richts, S., Fletcher, E.V., Vanwelden, T., Schmidt, B.Z., Henningfeld, K.A., et al. (2019). Prdm12 Directs Nociceptive Sensory Neuron Development by Regulating the Expression of the NGF Receptor TrkA. Cell Rep 26, 3522–3536 e3525.

Dong, X., and Dong, X. (2018). Peripheral and Central Mechanisms of Itch. Neuron 98, 482-494.

Dong, X., Han, S., Zylka, M.J., Simon, M.I., and Anderson, D.J. (2001). A diverse family of GPCRs expressed in specific subsets of nociceptive sensory neurons. Cell 106, 619–632.

Emrick, J.J., Mathur, A., Wei, J., Gracheva, E.O., Gronert, K., Rosenblum, M.D., and Julius, D. (2018). Tissue-specific contributions of Tmem79 to atopic dermatitis and mast cell-mediated histaminergic itch. Proc Natl Acad Sci U S A 115, E12091–E12100.

Fawcett, T. (2006). An introduction to ROC analysis. Pattern Recogn Lett 27, 861–874.

Fogh, K., Herlin, T., and Kragballe, K. (1989). Eicosanoids in skin of patients with atopic dermatitis: prostaglandin E2 and leukotriene B4 are present in biologically active concentrations. J Allergy Clin Immunol 83, 450–455.

Fontanet, P., Irala, D., Alsina, F.C., Paratcha, G., and Ledda, F. (2013). Pea3 transcription factor family members Etv4 and Etv5 mediate retrograde signaling and axonal growth of DRG sensory neurons in response to NGF. J Neurosci 33, 15940–15951.

Han, L., and Dong, X. (2014). Itch mechanisms and circuits. Annu Rev Biophys 43, 331–355.

Han, L., Ma, C., Liu, Q., Weng, H.J., Cui, Y., Tang, Z., Kim, Y., Nie, H., Qu, L., Patel, K.N., et al. (2013). A subpopulation of nociceptors specifically linked to itch. Nat Neurosci 16, 174–182.

Han, S.K., Mancino, V., and Simon, M.I. (2006). Phospholipase Cbeta 3 mediates the scratching response activated by the histamine H1 receptor on C-fiber nociceptive neurons. Neuron 52, 691–703.

Hu, G., Huang, K., Hu, Y., Du, G., Xue, Z., Zhu, X., and Fan, G. (2016). Single-cell RNA-seq reveals distinct injury responses in different types of DRG sensory neurons. Sci Rep 6, 31851.

Huang, C.C., Yang, W., Guo, C., Jiang, H., Li, F., Xiao, M., Davidson, S., Yu, G., Duan, B., Huang, T., et al. (2018a). Anatomical and functional dichotomy of ocular itch and pain. Nat Med 24, 1268–1276.

Huang, J., Polgar, E., Solinski, H.J., Mishra, S.K., Tseng, P.Y., Iwagaki, N., Boyle, K.A., Dickie, A.C., Kriegbaum, M.C., Wildner, H., et al. (2018b). Circuit dissection of the role of somatostatin in itch and pain. Nat Neurosci 21, 707–716.

Ikoma, A., Steinhoff, M., Stander, S., Yosipovitch, G., and Schmelz, M. (2006). The neurobiology of itch. Nat Rev Neurosci 7, 535–547.

Kathe, C., and Moon, L.D.F. (2018). RNA sequencing dataset describing transcriptional changes in cervical dorsal root ganglia after bilateral pyramidotomy and forelimb intramuscular gene therapy with an adeno-associated viral vector encoding human neurotrophin-3. Data in brief 21, 377–385.

Kido-Nakahara, M., Buddenkotte, J., Kempkes, C., Ikoma, A., Cevikbas, F., Akiyama, T., Nunes, F., Seeliger, S., Hasdemir, B., Mess, C., et al. (2014). Neural peptidase endothelin-converting enzyme 1 regulates endothelin 1-induced pruritus. J Clin Invest 124, 2683–2695.

Konishi, Y., Ikeda, K., Iwakura, Y., and Kawakami, K. (2006). Six1 and Six4 promote survival of sensory neurons during early trigeminal gangliogenesis. Brain Res 1116, 93–102.

Kremer, A.E., Martens, J.J., Kulik, W., Rueff, F., Kuiper, E.M., van Buuren, H.R., van Erpecum, K.J., Kondrackiene, J., Prieto, J., Rust, C., et al. (2010). Lysophosphatidic acid is a potential mediator of cholestatic pruritus. Gastroenterology 139, 1008-1018, 1018 e1001.

Kupari, J., Haring, M., Agirre, E., Castelo-Branco, G., and Ernfors, P. (2019). An Atlas of Vagal Sensory Neurons and Their Molecular Specialization. Cell Rep 27, 2508–2523 e2504.

Lallemend, F., and Ernfors, P. (2012). Molecular interactions underlying the specification of sensory neurons. Trends Neurosci 35, 373–381.

Langmead, B., and Salzberg, S.L. (2012). Fast gapped-read alignment with Bowtie 2. Nat Methods 9, 357–U354.

Leader, B., Carr, C.W., and Chen, S.C. (2015). Pruritus epidemiology and quality of life. Handb Exp Pharmacol 226, 15–38.

Li, C.L., Li, K.C., Wu, D., Chen, Y., Luo, H., Zhao, J.R., Wang, S.S., Sun, M.M., Lu, Y.J., Zhong, Y.Q., et al. (2016). Somatosensory neuron types identified by high-coverage single-cell RNA-sequencing and functional heterogeneity. Cell Res 26, 83–102.

Liu, Q., Sikand, P., Ma, C., Tang, Z., Han, L., Li, Z., Sun, S., LaMotte, R.H., and Dong, X. (2012). Mechanisms of itch evoked by beta-alanine. J Neurosci 32, 14532–14537.

Liu, Q., Tang, Z., Surdenikova, L., Kim, S., Patel, K.N., Kim, A., Ru, F., Guan, Y., Weng, H.J., Geng, Y., et al. (2009). Sensory neuron-specific GPCR Mrgprs are itch receptors mediating chloroquine-induced pruritus. Cell 139, 1353–1365.

Liu, Q., Weng, H.J., Patel, K.N., Tang, Z., Bai, H., Steinhoff, M., and Dong, X. (2011). The distinct roles of two GPCRs, MrgprC11 and PAR2, in itch and hyperalgesia. Sci Signal 4, ra45.

Lopez-Santiago, L.F., Pertin, M., Morisod, X., Chen, C., Hong, S., Wiley, J., Decosterd, I., and Isom, L.L. (2006). Sodium channel beta2 subunits regulate tetrodotoxin-sensitive sodium channels in small dorsal root ganglion neurons and modulate the response to pain.J Neurosci 26, 7984–7994.

Lou, S., Pan, X., Huang, T., Duan, B., Yang, F.C., Yang, J., Xiong, M., Liu, Y., and Ma, Q. (2015). Incoherent feed-forward regulatory loops control segregation of C-mechanoreceptors, nociceptors, and pruriceptors. J Neurosci 35, 5317–5329.

Love, M.I., Huber, W., and Anders, S. (2014). Moderated estimation of fold change and dispersion for RNA-seq data with DESeq2. Genome Biol 15, 550.

Meixiong, J., and Dong, X. (2017). Mas-Related G Protein-Coupled Receptors and the Biology of Itch Sensation. Annu Rev Genet 51, 103–121.

Meixiong, J., Vasavda, C., Green, D., Zheng, Q., Qi, L., Kwatra, S.G., Hamilton, J.P., Snyder, S.H., and Dong, X. (2019). Identification of a bilirubin receptor that may mediate a component of cholestatic itch. Elife 8.

Mishra, S.K., and Hoon, M.A. (2013). The cells and circuitry for itch responses in mice. Science 340, 968–971.

Mishra, S.K., and Hoon, M.A. (2015). Transmission of pruriceptive signals. Handb Exp Pharmacol 226, 151–162.

Mollanazar, N.K., Smith, P.K., and Yosipovitch, G. (2016). Mediators of Chronic Pruritus in Atopic Dermatitis: Getting the Itch Out? Clin Rev Allergy Immunol 51, 263–292.

Moody, S.A., and LaMantia, A.S. (2015). Transcriptional regulation of cranial sensory placode development. Curr Top Dev Biol 111, 301–350.

Morita, T., McClain, S.P., Batia, L.M., Pellegrino, M., Wilson, S.R., Kienzler, M.A., Lyman, K., Olsen, A.S., Wong, J.F., Stucky, C.L., et al. (2015). HTR7 Mediates Serotonergic Acute and Chronic Itch. Neuron 87, 124–138.

Oude Elferink, R.P., Bolier, R., and Beuers, U.H. (2015). Lysophosphatidic acid and signaling in sensory neurons. Biochim Biophys Acta 1851, 61–65.

Piomelli, D., and Sasso, O. (2014). Peripheral gating of pain signals by endogenous lipid mediators. Nat Neurosci 17, 164–174.

Qu, L., Fan, N., Ma, C., Wang, T., Han, L., Fu, K., Wang, Y., Shimada, S.G., Dong, X., and LaMotte, R.H. (2014). Enhanced excitability of MRGPRA3-and MRGPRD-positive nociceptors in a model of inflammatory itch and pain. Brain 137, 1039–1050.

Reddy, V.B., and Lerner, E.A. (2017). Activation of mas-related G-protein-coupled receptors by the house dust mite cysteine protease Der p1 provides a new mechanism linking allergy and inflammation. J Biol Chem 292, 17399–17406.

Reddy, V.B., Sun, S., Azimi, E., Elmariah, S.B., Dong, X., and Lerner, E.A. (2015). Redefining the concept of protease-activated receptors: cathepsin S evokes itch via activation of Mrgprs. Nat Commun 6, 7864.

Salvatierra, J., Diaz-Bustamante, M., Meixiong, J., Tierney, E., Dong, X., and Bosmans, F. (2018). A disease mutation reveals a role for NaV1.9 in acute itch. J Clin Invest 128, 5434–5447.

Sano, Y., Inamura, K., Miyake, A., Mochizuki, S., Kitada, C., Yokoi, H., Nozawa, K., Okada, H., Matsushime, H., and Furuichi, K. (2003). A novel two-pore domain K+ channel, TRESK, is localized in the spinal cord. J Biol Chem 278, 27406–27412.

Secundo, L., Snitz, K., and Sobel, N. (2014). The perceptual logic of smell. Curr Opin Neurobiol 25, 107–115.

Shim, W.S., Tak, M.H., Lee, M.H., Kim, M., Kim, M., Koo, J.Y., Lee, C.H., Kim, M., and Oh, U. (2007). TRPV1 mediates histamine-induced itching via the activation of phospholipase A2 and 12-lipoxygenase. J Neurosci 27, 2331–2337.

Solinski, H.J., Kriegbaum, M.C., Tseng, P.Y., Earnest, T.W., Gu, X., Barik, A., Chesler, A.T., and Hoon, M.A. (2019). Nppb Neurons Are Sensors of Mast Cell-Induced Itch. Cell Rep 26, 3561–3573 e3564.

Starobova, H., Mueller, A., Deuis, J.R., Carter, D.A., and Vetter, I. (2019). Inflammatory and neuropathic gene expression signatures of chemotherapy-induced neuropathy induced by vincristine, cisplatin and oxaliplatin in C57BL/6J mice. J Pain.

Storan, E.R., O’Gorman, S.M., McDonald, I.D., and Steinhoff, M. (2015). Role of cytokines and chemokines in itch. Handb Exp Pharmacol 226, 163–176.

Sullivan, S.J., Farrant, M., and Cull-Candy, S.G. (2017). TARP gamma-2 Is Required for Inflammation-Associated AMPA Receptor Plasticity within Lamina II of the Spinal Cord Dorsal Horn. J Neurosci 37, 6007–6020.

Suzuki, M. (2006). The Drosophila tweety family: molecular candidates for large-conductance Ca2+activated Cl-channels. Exp Physiol 91, 141–147.

Tao, F., Skinner, J., Su, Q., and Johns, R.A. (2006). New role for spinal Stargazin in alpha-amino-3-hydroxy-5-methyl-4-isoxazolepropionic acid receptor-mediated pain sensitization after inflammation. J Neurosci Res 84, 867–873.

Usoskin, D., Furlan, A., Islam, S., Abdo, H., Lonnerberg, P., Lou, D., Hjerling-Leffler, J., Haeggstrom, J., Kharchenko, O., Kharchenko, P.V., et al. (2015). Unbiased classification of sensory neuron types by largescale single-cell RNA sequencing. Nat Neurosci 18, 145–153.

Valtcheva, M.V., Samineni, V.K., Golden, J.P., Gereau, R.W.t., and Davidson, S. (2015). Enhanced nonpeptidergic intraepidermal fiber density and an expanded subset of chloroquine-responsive trigeminal neurons in a mouse model of dry skin itch. J Pain 16, 346–356.

Wan, L., Jin, H., Liu, X.Y., Jeffry, J., Barry, D.M., Shen, K.F., Peng, J.H., Liu, X.T., Jin, J.H., Sun, Y., et al. (2017). Distinct roles of NMB and GRP in itch transmission. Sci Rep 7, 15466.

Waxman, S.G., and Zamponi, G.W. (2014). Regulating excitability of peripheral afferents: emerging ion channel targets. Nat Neurosci 17, 153–163.

Wilson, S.R., Gerhold, K.A., Bifolck-Fisher, A., Liu, Q., Patel, K.N., Dong, X., and Bautista, D.M. (2011). TRPA1 is required for histamine-independent, Mas-related G protein-coupled receptor-mediated itch. Nat Neurosci 14, 595–602.

Yajima, H., Suzuki, M., Ochi, H., Ikeda, K., Sato, S., Yamamura, K., Ogino, H., Ueno, N., and Kawakami, K. (2014). Six1 is a key regulator of the developmental and evolutionary architecture of sensory neurons in craniates. BMC Biol 12, 40.

Yarmolinsky, D.A., Zuker, C.S., and Ryba, N.J. (2009). Common sense about taste: from mammals to insects. Cell 139, 234–244.

Yosipovitch, G., and Bernhard, J.D. (2013). Clinical practice. Chronic pruritus. The New England journal of medicine 368, 1625–1634.

Zhu, Y., Hanson, C.E., Liu, Q., and Han, L. (2017). Mrgprs activation is required for chronic itch conditions in mice. Itch (Phila) 2.

Zimmerman, A., Bai, L., and Ginty, D.D. (2014). The gentle touch receptors of mammalian skin. Science 346, 950–954.

